# Female-dominated disciplines have lower evaluated research quality and funding success rates, for men and women

**DOI:** 10.1101/2024.03.14.585000

**Authors:** Alex James, Franca Buelow, Liam Gibson, Ann Brower

## Abstract

We use data from 30 countries and find that the more women in a discipline, the lower quality the research in that discipline is evaluated to be and the lower the funding success rate is. This affects men and women, and is robust to age, number of research outputs, and bibliometric measures where such data are available. Our work builds on others’ findings that women’s work is valued less, regardless of who performs that work.

## Introduction

Many have observed that women’s research is evaluated as lower quality than men’s research. [1, 2] And apart from a few exceptions [3–5], most analyses show that women’s funding success rates are lower than men’s [6–8] both in terms of number of grants and size of grants [7, 9]. Evaluations of research quality affect many aspects of an academic career, from hiring and promotion decisions to funding success. Other studies have explored possible explanations for this seemingly gendered pattern in evaluation of research quality, such as societal [1, 8, 10], institutional [11], and academic factors [12, 13], including gender bias [14].

We find a strong correlation between the evaluated quality of a researcher’s work and the gender balance of the researcher’s discipline. Researchers, both male and female, in female-dominated disciplines receive lower evaluations. This finding is robust to bibliometric differences across disciplines. We also find that funding success rates display a similar correlation. Applications from researchers, both male and female, in female-dominated disciplines have lower success rates.

Academic disciplines can be quite different to one another in ways such as publication rates[15], citation rates [16] and expected h-indices [17]. There are different norms for the expected number of co-authors[18], order of authors, and attributions [19]. Gender balance is not constant across disciplines either. There are fewer women in STEM subjects, and more women than men use qualitative methods, rather than quantitative, in research [14]. Recent research suggests there are interactions between individual and discipline level characteristics. Higher ‘expectations of brilliance’ in a discipline is related to a lower proportion of women entering [20]; and research area is a strong contributor to the lower funding success rates of African American/Black scientists [21].

That women score lower on evaluations of research quality is not surprising. On an individual level, women publish less than men [2, 22, 23], receive fewer citations [24, 25], and have lower h- indices [26]. Women have fewer first authorships [2] and are less likely to receive acknowledgment [27]. Women are also less likely to get credit for their innovations [28, 29] and awards, particularly prestigious ones, are less likely to be given to women [30, 31]. Recent research suggests women react differently than men to a change in a journal’s prestige or impact factor [32]. However, our research shows a different correlation: a higher proportion of women in a discipline is correlated with lower research quality evaluations and funding success rates, for everyone in the field.

We use independent datasets spanning thousands of researchers and 30 countries and look separately at research funding success rates and holistic research quality scores. Where possible we account for researcher characteristics like age, research institution and publishing patterns. We consistently find that researchers in male-dominated disciplines have both higher funding success rates, and higher research quality scores, than researchers in female-dominated disciplines. This applies to both men and women.

## Data and Methods

We analysed research quality and funding success separately. We used data from 4 sources (PBRF, ARC, CIHR, EIGE), spanning 30 countries (described fully in Supplementary Materials). The ARC, CIHR and EIGE datasets cover funding success rates across 29 countries. The PBRF dataset covers research quality evaluations for all university academic staff in New Zealand. Summary data from the 4 datasets are reported in Table 1. The discipline category definitions varied in each dataset, e.g. EIGE data used Science as a discipline whereas PBRF broke this down into smaller groupings like physics, biology etc. Supplementary material contains full information for each dataset. Due to data limitations, we consider gender as binary, data on non—binary individuals were either not available (EIGE and CIHR) or the group was too small to be included (PBRF and ARC).

**Table 1:** A summary of the four independent datasets used in the study showing: country, time period of the data, number of disciplines, research evaluation method, number of individuals or applicants by gender, number of individual data points, output variable and mean expected output for each gender across the whole dataset.

| Scheme | PBRF |  |  | ARC | CIHR | EIGE |
| --- | --- | --- | --- | --- | --- | --- |
| Country | Aotearoa, New Zealand |  |  | Australia | Canada | Predominantly EU |
| Timespan | 2000-06 | 2007-12 | 2013-18 | 2010-19 | 2011 - 16 | 2019 |
| Disciplines | 42 | 42 | 43 | 22 | 4 | 8 |
| Evaluation | Research Quality Score (200 – 700) |  |  | Funding success |  |  |
| Women | 1708 | 2658 | 3297 | 46,231 | 8,143 | 49,863 |
| Men | 2522 | 4005 | 4181 | 130,331 | 15,775 | 85,305 |
| Data points | 4230 | 6663 | 7487 | 440 | 16 | 333 |
| Outcome | Mean score |  |  | Success rate |  |  |
| Women | 272 | 368 | 379 | 25.7% | 14.7% | 27.0% |
| Men | 351 | 424 | 433 | 27.1% | 16.4% | 30.3% |

*Research quality evaluations:* we use the Aotearoa, New Zealand (ANZ) Performance Based Research Fund (PBRF) data. This dataset contains holistic research portfolio evaluations of every research active academic in ANZ over three time spans (2000-06, 2007-12, 2013-18). The holistic research portfolio includes an individual’s self-nominated best three or four publications, in full, with descriptions of the importance and impact of the publications. It also includes non-publication aspects such as contributions to the research environment like chairing a conference organisation committee. Evaluation is done by subject area panels of experts, who read the publications and evaluate the individual’s holistic contribution. It is not an automated process and does not explicitly use measures such as h-indices, number of publications or number of citations. The evaluation explicitly claims to focus on quality over quantity [33]. Panels are likely to be more gender balanced than the fields they assess but in the more heavily skewed fields (e.g. Education, Engineering, Mathematics) they are skewed as would be expected. The PBRF is described in more detail in Supplementary Material.

With over 18,000 individual data points this dataset is a unique opportunity to explore a question such as this. To examine the robustness of any findings to bibliometric measures, we use a subset of the PBRF data from one university to compare anonymised detailed research output data to each individual’s PBRF score.

*Funding Success Rates:* we use three independent publicly available datasets: Australian Research Council (ARC) application success rates from 2010 to 2019; Canadian Institute of Health Research application data from 2013 to 2016; and data covering government research grants across the EU and UK in 2019 from the European Institute of Gender Equality (EIGE). These data sets all provided aggregated data on the number of applications and successes by gender for each research discipline, i.e. not individual level data.

The data sets are briefly described in the text, Table 1 and (Table S1) and full data details are in Supplementary Materials. Each dataset is analysed separately and the analysis includes the sample size of each aggregated datapoint. We define a maximal model including two-way interactions between the three key variables: the gender balance of the discipline measured by the proportion of researchers who are men, *p*_men_; researcher gender, *Gender*; and, where appropriate, *Time*. Any other available variables, e.g. researcher age, institute or country were included as linear terms with either a fixed (Age) or random (Institute or Country) effect. We measure ‘best’ by the lowest AIC and the choice is robust by other similar measures e.g. maximising Pearson’s R-squared or omitting non-significant variables. Table 2 shows the maximal models and the best models for each of the datasets.

**Table 2:**
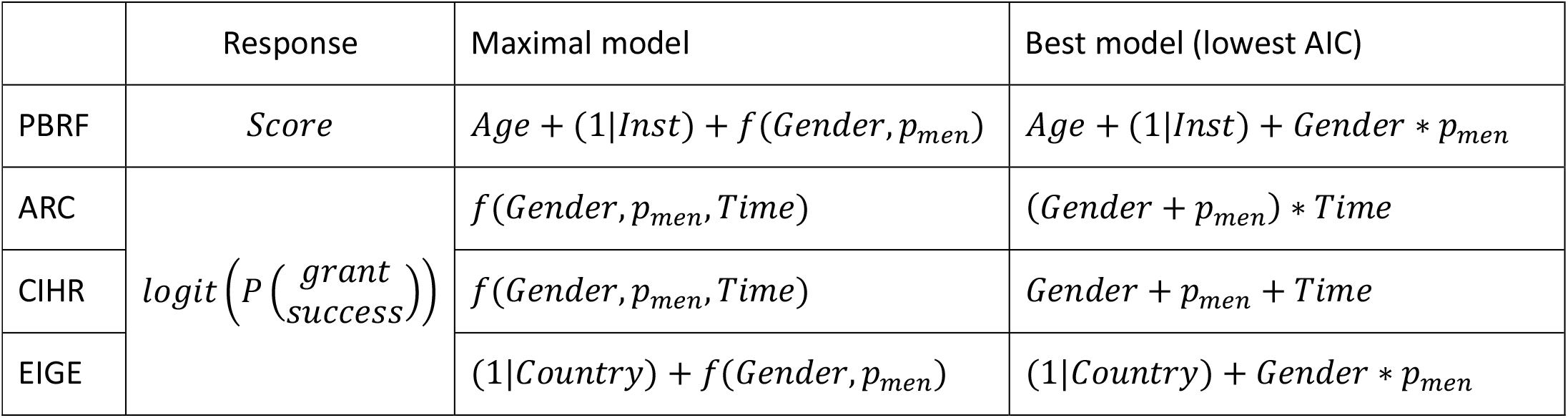
Maximal candidate model and best fit model to predict *Score* for PBRF and *logit*(*P*(*Success*)) for each dataset. Wilkinson notation is used to indicate interaction terms.

### Scoring research quality

First, we look at the only country that scores all individual university researchers. The Aotearoa New Zealand Performance-Based Research Fund (PBRF) scores are individual level micro-data on the results of 18,371 holistic research quality assessment scores of 13,555 individuals in ANZ for the PBRF assessments. The assessments were carried out at three separate time points. In each of 2006, 2012, and 2018, staff members at all of the eight universities in ANZ, were given a research performance score from 0 to 700 based on a holistic assessment of their research portfolio over the previous 6 years. Individuals with scores over 200 were considered research active. At each assessment, between 4,000 and 8,000 researchers were assessed. The scores, based on assessments by local and international experts on 14 different subject area panels, were used to allocate government performance-based research funding to universities. Although many researchers were present for more than one round, we analyse each of the three time points separately to examine the link between research score and the gender balance of a researcher’s discipline.

We use a linear regression model to predict the research quality score using the available variables, in particular the gender balance of the discipline, *p*_men_. Although the data is strictly count data requiring a Poisson model we use a simple linear model as all the predicted values are large, i.e. between 200 and 700. We start with the maximal model of Table 2 that includes our key variables of interest *Gender* and *p*_men_ with interactions and the confounding variables *Age* and *Institute* as linear terms with no interactions. We compare all candidate models up to the maximal model and choose the best model by minimising AIC. The electronic Table E2 gives the full output from all candidate models. For all datasets all terms in the best model (PBRF3 in TableE2) were significant (*p* < 0.05) and the best model had a Δ*AIC* > 2 compared to the second best.

The best model for all three time points was

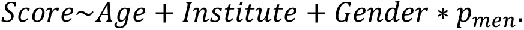

The model predicts that individuals in disciplines dominated by men have higher research performance scores than individuals in female-dominated disciplines; all variables are significant (*p* < 0.05, see Table E2) at all time points.

Even after accounting for age in this way, men have higher scores than women; but the difference between the most male-dominated disciplines and the most female-dominated is larger than the gap between men and women within any particular discipline. Figure 1 shows the mean raw score of each discipline by gender (women red +, men blue x) in each assessment period (A) 2006, (B) 2012, (C) 2018. The line is the expected model output for a 50-year-old male (blue line) or female (red line) researcher at the University of Canterbury (UC) (other ages and institutions differ by a constant as they are included as linear terms with no interactions). Note that the full model predictions (lines on Fig 1) are adjusted for age so may appear different from the raw data (points on Fig 1). In particular, raw data points from discipline/gender groups with many young researchers will appear much lower than might be expected at first thought.

**Figure 1:**
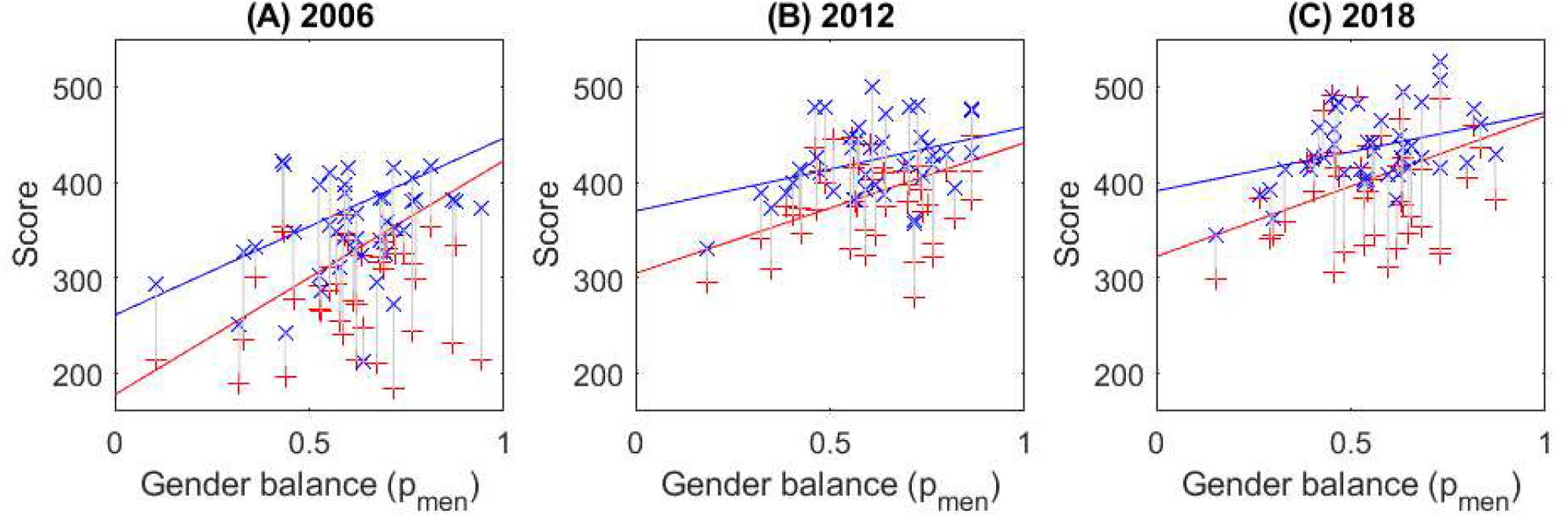
Male-dominated disciplines have higher expected research scores than female-dominated disciplines in all three PBRF assessments. Points – the mean raw score of individuals in each of the disciplines (full raw data cannot be shown for privacy reasons), against the proportion of men in the discipline (blue – men, red – women). Lines – expected score from individual level analysis (*N* = 4135, 6586, 7467 respectively) adjusting for age, institute, gender, and proportion of men in the discipline, and shown for a 50-year-old man (blue) and woman (red) at one university in Aotearoa NZ.

By way of example, we consider a 50-year old researcher in 2018 at University of Canterbury (i.e. lines in Figure 1C), *Institute* and *Age* are linear terms in the model so a different aged researcher at another university would show a very similar pattern. We assume our researcher comes from either a male-dominated discipline (*p*_men_ = 75%) like Philosophy or Physics; a gender balanced discipline (*p*_men_ = 50%) such as Psychology or Law; or a female-dominated discipline (*p*_men_ = 25%) like Nursing or Education., In the gender-balanced discipline (Psychology or Law) a woman’s expected score is 396 out of a possible 700. This is around 8% lower than a man’s expected score of 432/700 in the same discipline. If a man works in a male-dominated discipline (Physics or Philosophy), his mean score is now expected to be higher at 452/700. If a man works in a female-dominated discipline (Nursing or Education), his score will drop to 411/700. In comparison, a female Philosopher or Physicist can expect a score of 433/700, only slightly lower than a male researcher in the same discipline. If a woman works in Nursing or Education, her expected score is now 359/700, a drop of over 20% from the female Philosopher’s expected score and also lower than the male Nursing researcher’s score.

This effect, of researchers in female-dominated disciplines being predicted to have lower scores than researchers in male-dominated disciplines, persists for all three assessment rounds but was much larger in the 2006 round (Fig 1A). In all three rounds, the score drop associated with moving from a male- to a female-dominated discipline is larger for women than for men. Finally the gender gap within a discipline is much larger in female-dominated disciplines than in male–dominated ones. Though the gap has closed considerably between 2006 and 2018.

### Accounting for bibliometric differences between individuals

Here we explore whether gendered patterns in publication explain the variation in research scores across disciplines. We show that publications are important but they do not negate our previous result that researchers in female-dominated disciplines receive lower research scores.

We analysed a subset of data available from UC, one of the eight universities contained in the PBRF datasets. We included research-active staff, (i.e. PBRF Score > 200) with at least one publication between 2013-18, working at UC in 2018. Using date of birth, gender, ethnicity, and discipline we were able to link the university’s internal research publications dataset to individual PBRF scores for over 80% of individuals (*N* = 384, 141 female, 243 male). Data records that could not be linked were excluded.

These publications records allowed us to calculate a series of research bibliometrics e.g. number of outputs, number of citations, expected field weighted citation index of publications etc, for each researcher. We then used these bibliometrics to predict each individual’s expected research score (see Supplementary Material for a more detailed description of the data and the analysis). The bibliometrics were tested separately in an individual regression model

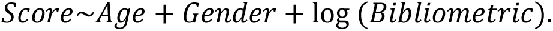

The bibliometric that was the best predictor of research score (with minimum AIC and maximum r-squared) was the total number of outputs *N*_outputs_ (See Table E3, model 2). This output gave a model correlation of *r*^2^ = 0.325 and the coefficient of *N*_outputs_ was highly significant (*p* < 0.001), In this model *Age* was also significant (*p* < 0.01) but *Gender* was not (*p* > 0.05). The second-best bibliometric predictor by AIC, number of outputs weighted by number of authors was also a measure of research quantity, It had correlation *r*^2^ = 0.280. By comparison the other bibliometrics, all measures more related to research quality, had *r*^2^ < 0.09. We also tested a two bibliometric model by combining the best predictor, *N*_outputs_, with each of the other bibliometrics separately. In all cases the change in AIC due to the addition of the second bibliometric was very small (Δ*AIC* < 2) and the secondary bibliometric was not significant (*p* > 0.05) (see Table E2).

The bibliometric analysis shows several things. First, Figure 2A shows the best available predictor of research score was number of research outputs; and bibliometrics related to quality (e.g. expected field weighted citation index) had almost no effect. Alone, this result is surprising as documentation on PBRF clearly states that the evaluations are based on quality not quantity of research [33]. Similarly, *Gender* is not a predictor of research quality, a man and a women of the same age with the same number of research outputs have the same expected score. Figure 2A shows the raw scores of all individuals in the single university sample and the predicted relationship between number of outputs and Score for a 50-year-old of either gender. On average, men have more outputs than women (Figure 2A vertical error bars) and get higher scores (horizontal error bars).

**Figure 2:**
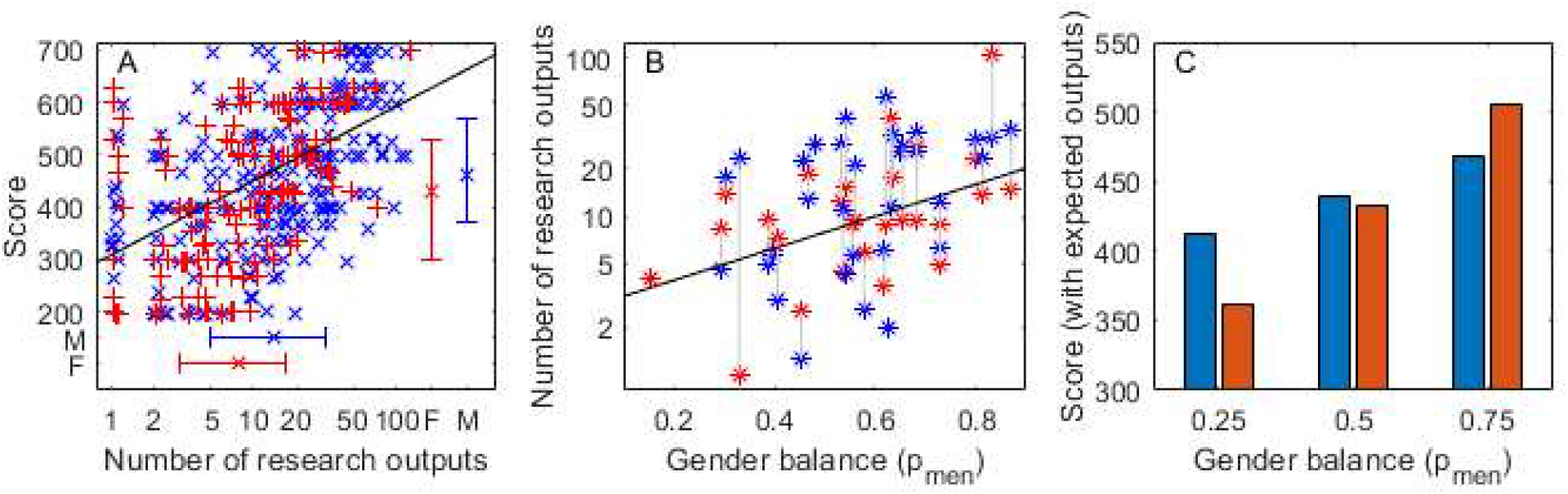
Research score is strongly correlated with number of research outputs (2A), which is correlated to the gender balance of the discipline (2B); but, after adjusting for a typical number of research outputs, female-dominated disciplines still have lower scores (2C). PBRF data using individuals at UC (N=384). A) Score against number of research outputs over the period of the PBRF assessment. Trend line shows the expected score for a 50-year old. Error bars show the median and interquartile range for men and women for score (vertical) and number of outputs (horizontal). B) Mean number of outputs by gender for each discipline against gender balance of the discipline. Trend line from individual level analysis (*N* = 384), gender was not significant. C) Expected score of a 50-year-old man and women with the expected number of outputs for a discipline of that gender balance (25%, 50%, 75% men respectively).

Having seen that score is strongly correlated to the number of research outputs (Fig 2A), we then test the relationship between the number of outputs and the proportion of men in the discipline using the same sample of 384 individuals at UC (Fig 2B). We start with the maximal model to allow the relationship to change for each gender

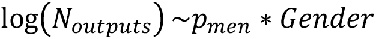

The gender balance of the discipline is highly significant (coefficient *p*_men_: *p* < 0.001) but again *Gender* is not significant either alone or with an interaction (*p* > 0.05). So the most parsimonious model is

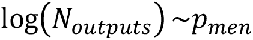

meaning a man and woman in the same discipline have, on average, the same number of outputs and researchers in male-dominated disciplines have, on average, more publications. Taken together, it is tempting to use these correlations as an explanation for the lower scores for researchers in female-dominated disciplines. In other words, both men and women in female-dominated disciplines publish less, and fewer publications lead to a lower score. This explanation fails the empirical test because, as seen in Figure 1, on average a man receives a higher score than a woman in the same discipline. This is *despite*, on average, having the same number of outputs (Figure 2B).

Finally, we test the relationship of our previous analysis relating *Score* to *Gender* and gender balance, *p*_men_, but now we add the best bibliometric predictor *N*_outputs_

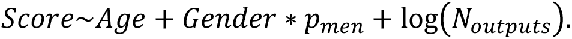

After using this model to account for the difference in publication norms across disciplines with different gender balance, the proportion of men in the discipline is still a significant predictor of score (coefficient and interaction *p* < 0.05, see Table E3), but now the relationship is more intricate.

In a female-dominated (*p*_men_ = 25%) discipline, like Nursing or Education, the expected number of outputs is low for both genders (Fig 2B) and scores are correspondingly low. But given that a man and a woman have the same expected number of outputs for their discipline (as predicted by Fig 2B), the man’s expected score of 412/700 is higher than the woman’s of 361/700 (Fig 2C, left bars). When we consider a male-dominated (*p*_men_ = 75%) discipline, like Physics or Philosophy, given gender-equal (but now likely to be high) publication rates, overall scores are much higher, a man’s score is predicted to be 467/700 and a woman’s is higher still at 505/700 (Fig 2C, right bars). In short, even after accounting for variations in publication rates, researchers in female-dominated disciplines still have lower expected research scores.

In this detailed look at PBRF scores compared to bibliometric measures, the gender minority has an advantage over the majority at both ends of the gender balance spectrum. If those in the gender minority of a discipline have the same number of publications as their majority colleagues (as our findings suggest they usually do), the gender minority will, on average, receive a higher research evaluation score.

Here we have presented a more detailed analysis which accounts for publication differences. Whilst differences between the gender minority and majority in a discipline continue to exist, the key point remains: when women are in the majority, the scores are lower for everyone (Figure 2C).

Far more men work in male-dominated disciplines (PBRF 2018: 74% of men in disciplines with *p*_men_ > 50%) than women (PBRF 2018: 47% of women in disciplines with *p*_men_ > 50%). Hence men will benefit overall from a pattern of evaluating research in male-dominated disciplines as higher quality than research in female-dominated disciplines.

## Research funding success rates

We examine funding success rates using three independent datasets. Table 1 describes the datasets and shows summary statistics including the number of individuals each data set covers and the resulting sample size. The raw data in Table 1 suggest women have lower success rates than men across all the datasets.

We use three independent datasets from: the Australian Research Council (ARC), the Canadian Institute of Health Research (CIHR) and the European Institute of Gender Equality (EIGE). Each dataset gives aggregated data on the number of applicants and the number of successes, by gender and discipline, for research funding. For the ARC and CIHR this is over a number of years, for EIGE this is in a single year. For EIGE the gender balance of each discipline was available separately for over 80% of the dataset and covered all researchers in the discipline nationally, in the remaining cases it was approximated with the applicant gender balance. For ARC gender balance was available separately (see Supp Mat for details) and again covered the whole population though it was only for a single year (2018). Data is also available for 2015 but as no discipline changed by more than 2 percentage points between 2015 and 2018 it was not used here. For CIHR we used an approximation based on the number of applicants.

We analyse each dataset separately and use a logistic regression model to predict the funding success rate. As our three datasets vary considerably rather than choose a model *a priori* we start with a maximal model that includes the key variables *Gender*, *p*_men_ and, when available, *Time* with up to two-way interactions between all variables. Any additional available variables are included as linear terms (see Table 2). We compare all candidate models up to the maximal model and choose by minimising AIC. Analysis was done on raw data so each group was weighted by sample size. Detailed data descriptions are given in Supplementary Materials.

### Australian Research Council (ARC)

The data included applicant numbers and successes across 22 disciplines, corresponding to the Australian Fields of Research classifications, over the 10 years from 2010 to 2019. See Supplementary Material for a more detailed data description. Full output from all models tested is in Table E2 and data are available supplementary material. The best fit model was

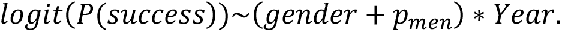

All predictors and interactions are highly significant (*p* < 0.001, see Model ARC2 Table E2).

In 2010, there was a fairly constant disadvantage for women. Gender-balance of the discipline had a relatively minimal effect on success; for example, in female-dominated (*p*_men_ = 25%) disciplines like Nursing a man’s chance of success was 32.6% and hers 30.1%. Similarly, in a male-dominated discipline (*p*_men_ = 75%) like Philosophy a man’s success rate would have increased marginally to 33.1% and a woman’s to 30.5% (solid lines, Figure 3A). In 2010, women’s disadvantage in grant applications was affected more by their gender than by their discipline.

**Figure 3:**
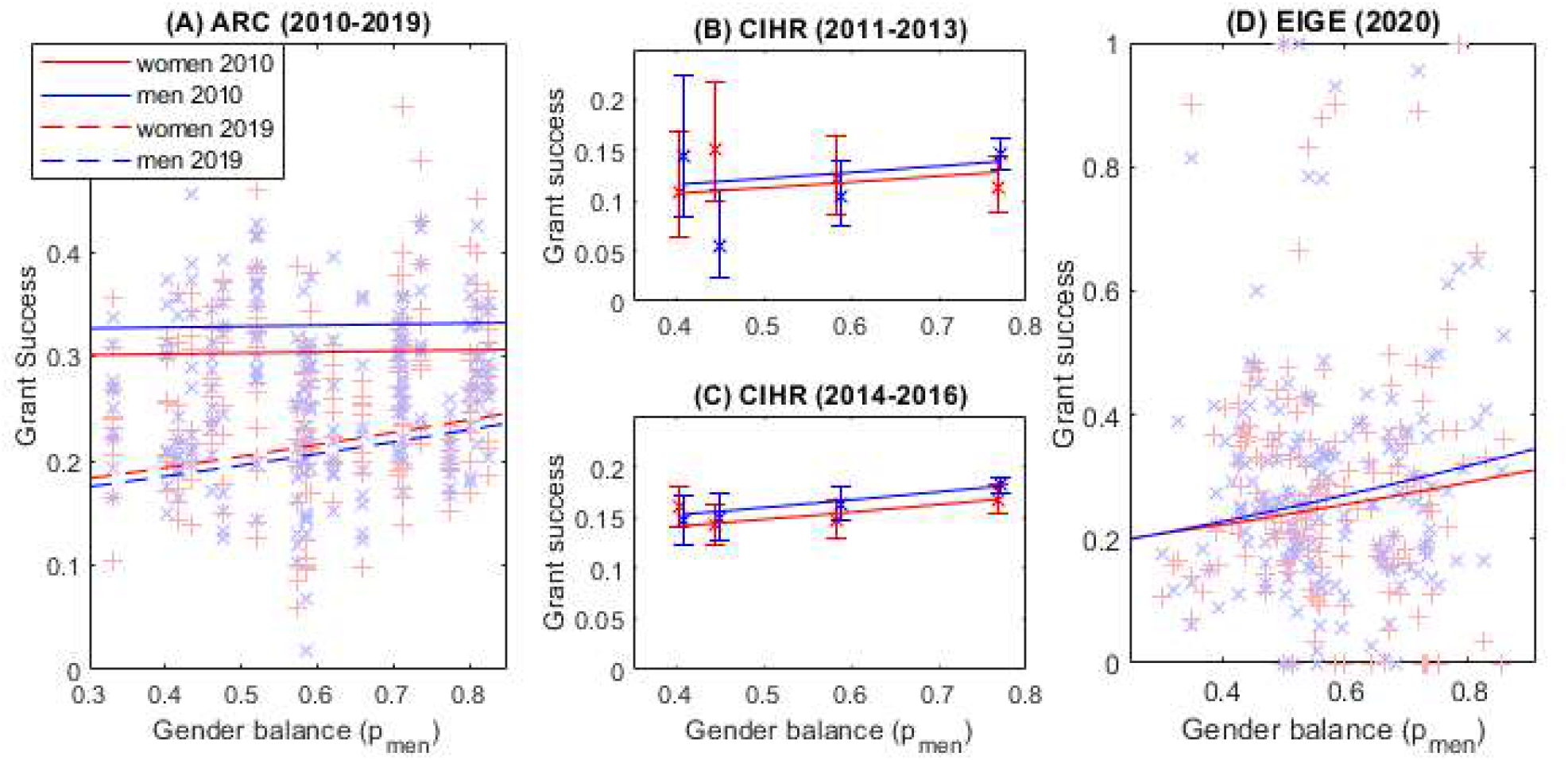
Researchers in male-dominated disciplines have a higher chance of funding success. A) ARC (2010-2019): Points - funding success rates by gender in 20 disciplines over 10 years against proportion of men in the discipline in 2018. Lines – expected success rates of men and women in 2010 (solid lines) and 2019 (dashed lines). B) CIHR (2011-2013), C) CIHR (2014-2016): Points – funding success rates by gender for each discipline against proportion of men in the discipline (estimated from application numbers) error bars are binomial 95% CI. Lines – expected success rate from combined analysis of all grant types in both time periods. D) EIGE (2020): Points - funding success rates by gender in 8 disciplines and 27 countries against proportion of men in the discipline in that country. Lines – expected success rates of men and women.

By 2019, overall success rates were much lower (dashed lines Figure 3A). The effect of gender was now reversed, giving marginally higher success rates to women but only when compared to men in the same discipline. In other words, a male Psychology researcher (*p*_men_ = 50%) has a 19.6% success rate compared to a female researcher in the same field at 20.4%. But now the discipline’s gender balance has a much larger effect. A male Philosopher’s (*p*_men_ = 75%) success rate has increased to 22.4% and a female Philosopher’s success rate is even higher at 23.3%. However, in Nursing (*p*_men_ = 25%), a female applicant has only a 17.8% probability of success; this is slightly higher than the male applicant’s 17.0% probability of success. Compared to 2010, the gender disadvantage has been replaced by the larger disadvantage of being in a female-dominated discipline.

### Canadian Institute of Health Research (CIHR)

This dataset was published in [6] and includes the number of applications and success rates by gender from 23,000 applicants to the CIHR in four sub-disciplines of Medical Science (Clinical, Biomolecular, Public Health, and Health Sciences). However, this is aggregated data so the number of datapoints is much smaller (see Supplementary Material Table S1). The time period used was before and after a change in funding model in 2013. See Supplementary Material for a more detailed data description. Full output from all models tested is in Table E2 and data is available in Supplementary Material. The best model (model CIHR2 Table E2) was

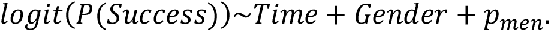

The original study, which accounted for age, found the discipline with the highest proportion of men (Biomedical, 77% men) had significantly higher odds of success than the three other disciplines, with 41% − 59% men Witteman, Hendricks (6) at Tables 2 and 3]. We do not have the age data controlled for in the original study, but we do find that there is a significant relationship (*p* < 0.001) between grant success and the proportion of men in the discipline. Unlike the ARC data men are slightly more likely to be successful overall. In both time periods Canadian health sub-disciplines that are more male-dominated have a higher chance of funding success (before the funding change - Fig 3B, after the funding change – Fig 3C). Our model predicts that the advantage of working in the most male dominated discipline (Biomedical 77% male) compared to the most female dominated (Public Health, 41% men) is almost three times the size of the direct advantage of being male.

### European Institute of Gender Equality (EIGE)

This dataset contains the number of applications and success rates by gender and country of origin from over 135,000 applicants to Government research funds predominantly in the EU but also including the UK, Israel and Turkey in 2019. Applications are divided into broad research areas, e.g. Science, Engineering, Humanities etc. Data was not available for all 8 disciplines in every country. See Supplementary Material for a more detailed data description. Full output from all models tested is in Table E2 and data is available in Supplementary Material. The best model (model EIGE3 Table E2) was

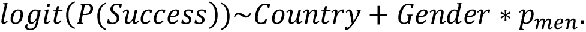

All variables are significant (*p* < 0.05). In male-dominated fields women have a slightly lower success rate in male dominated disciplines and, as with the other funding data sets, there is an overall increase in success rates as the discipline becomes more male-dominated. Once again, the effect on success rate of the applicant’s gender is small compared to the effect of the gender balance of the applicant’s discipline (Figure 3D). In the most male dominated disciplines (80% male) a man has a two percentage point advantage over female colleagues; by comparison a man has an 11 percentage point advantage over all researchers working in the most female dominated discipline (20% female). In relative terms this results in researchers in male dominated disciplines being 50% more likely to be successful in funding applications than researchers in male dominated disciplines.

## Discussion

We present two similar, yet separate, findings: 1) gender balance in a discipline correlates with research quality scores; and 2) gender balance in a discipline correlates with research funding success rates. These findings do not identify a causal relationship in which gender balance causes low research scores or funding success or vice-versa. But we can highlight and explore some possible explanations and theories that might link gender balance and research score/funding success. Further, even without establishing causality, these paired findings offer another piece of the academic gender puzzle.

### Feminization

Some have found that as a workforce feminises, salaries fall [34]. To explore whether as a discipline feminised the evaluation of research quality in that discipline also changed, we would need extensive longitudinal data. We cannot do that with the available data. But it is possible that the finding that research in female-dominated disciplines is evaluated as less good than research in male-dominated disciplines is an outcome of feminisation over time.

### Individual Gender Bias

One possible explanation of our findings is simply that reviewers (of research portfolios in NZ, and of research proposals in Australia, Canada, and the EU and UK) are biased against women. Indeed the effects of unconscious bias on academics are theorised thoroughly [35, 36]. Although empirics are limited [37] it has been seen that unconscious biases can affect common academic metrics and practices such as h-index, citation, authorship, and peer-review, as well as hiring and progression [38].

The PBRF datasets offer the most granular data and hence the best opportunity to explore gender bias on an individual level. For example, if the PBRF was dominated by individual bias this would manifest itself in Education having lower average scores than Physics but a woman in Education would have the same score as a woman in Physics on average. We see the former, but not the latter. However we do see that across all disciplines women have lower scores than men and this is more pronounced in female-dominated disciplines which is suggestive of gender bias but far from conclusive due to the many factors at play.

Further, the bibliometric subnational dataset suggests that a man and a woman with the same number of outputs (the best bibliometric predictor of score) have the same expected score Figure 2B). But when we include discipline gender bias (Figure 2C) the picture is less clear suggesting that if we compare a man and a woman with the same number of outputs the person in the minority gender is likely to have a higher score.

### Women do lower quality research

A small elephant in the room that might explain our findings is simply that women do lower quality research than men. Our findings do not show this. In 2018 in a male-dominated subject like Physics, a woman’s score is expected to be the same as a man’s (Figure 1). Similarly, an Australian woman has a slightly higher funding rate than a man in the same discipline in the ARC. These point to men and women having very similar research quality after accounting for discipline.

We do see that researchers, both male and female, in female-dominated disciplines have fewer research outputs on average. This could be due to cultural norms in these disciplines being driven by women who overall have fewer outputs; or it could be that women are attracted to disciplines with lower publication norms.

### Pre-allocation of and quotas for research funds

For research funding datasets (ARC, CIHR, EIGE), quotas or pre-allocation of funds towards certain types of research (e.g. public health funding increasing during the Covid pandemic) could explain some of our findings. But these pre- allocations would need to be towards male-dominated disciplines. Such a gendered pattern of pre-allocation suggests it would be more of a different mechanism for our findings, rather than an explanation of them. But we examine the datasets in turn against this explanation.

Most ARC applications are to the Discovery Project fund. Here, applications are scored independently and there are no pre-allocation decisions to value or fund one discipline over another [31]. Application success rates in 2023 are similar across the five very broad fields of research. This supports ARC’s claim that the process does not overtly pre-allocate by discipline. However, when we consider applicant success rates over the more detailed 22 disciplines, female-dominated disciplines have lower success rates.

There is little detailed data about the underlying processes that lie behind the EIGE data. Every country has its own internal rules and priorities for funding allocation. If local policy prioritises funding to Engineering and Physical Science, this could result in higher funding success rates in these areas; although it could also result in these areas having more researchers and the success rate being unaffected. Similarly we have no information on the Canadian funding system.

Conversely, in PBRF, this effect of pre-allocation is negligible or even non-existent. Every individual is scored independently. A surfeit of high scoring individuals in Physics does not stop high scores in Education. Overall, the pre-allocation of funds cannot explain all our results; and our results appear robust to this explanation.

### Job choice

This is frequently offered as an explanation of the gender pay gap. Women choose to work in job fields that, coincidently, are paid less [39]. It is a possibility that discipline choice, as opposed to job choice, is at play here, where women choose to work in disciplines where research is seen as less good. In academia, women tend to choose research areas and academic disciplines that the educational system undervalues [40]. Additionally, within a discipline, women tend to choose research topics that are more interdisciplinary and more applied [41].

Several individual and contextual factors affect job choice and evaluations [42]. Perceptions of excellence can correspond with an underrepresentation of women in certain disciplines, resulting in a higher perceived genius status for that discipline [20]. Our results find a correlation but not the causation for female-dominated disciplines having lower evaluations of research quality. Hence we can neither support nor reject an explanation of job choice.

### Bias against female-dominated disciplines

A simple explanation for our findings is that evaluations of research are biased but not against individuals as considered above earlier. Rather this explanation posits a bias against research disciplines dominated by women. Again as our findings are correlation not causation, we cannot accept or reject this explanation with certainty. Instead we can point to other research that suggests that we perceive male-dominated disciplines to require “brilliance”, while female-dominated disciplines require “hard work” [20]. If these discipline specific perceptions spill over into evaluations of research and funding applications, it could produce a gender-based discipline bias very similar to our findings.

Another simple explanation is that research in female-dominated disciplines is of lower quality than research in male-dominated disciplines. Research varies widely across disciplines from the quantitative “hard sciences” to the qualitative “soft sciences” but to categorize the efforts of whole disciplines as lower quality rather than different is an extreme step. This explanation also raises the question of why disciplines perceived as lower quality are more likely to be female-dominated.

Speculation aside, our findings do not show that research done in disciplines dominated by women is of lower quality. They do suggest that we consistently evaluate research from female-dominated disciplines as lower quality than research from male-dominated disciplines. This has a detrimental effect on everyone in those disciplines, but will adversely affect more women than men.

These findings will have implications in labour environments where research funding or research score affect salary or rank of individuals [1]. Future research could include modelling of various “levers of change” [11] that might serve to diminish, correct, or compensate for the gendered patterns observed.

Our datasets do have limitations. They cover a range of countries but are not representative of all of academia worldwide. And neither research evaluation nor funding success is a perfect measure of academic quality.

## Conclusion

We present two separate findings: gender balance of the discipline is strongly related to research quality evaluations and to funding success. We cannot claim to establish causality nor combine the two findings. Instead we seek possible explanations for our findings in others’ work.

Our findings suggest that what is perceived as women’s research is valued less, whether it is a man or a woman doing the research and whether or not overt bias is to blame. Regardless of causality, our findings suggest that patterns of devaluing women’s work affect all who do it, regardless of gender. And our findings point to areas that are ripe for further exploration.

## Supporting information

DataCIHR

DataEIGE

DataARC

## Acknowledgements

We thank Pen Holland, Elena Moltchanova and Ximena Nelson for providing comments on a draft of this work.

## Author contributions

Conceptualization: AJ, AB, FB

Methodology: AJ, AB, FB

Analysis: AJ

Investigation: AJ, AB, FB, LG

Writing – original draft, review and editing: AJ, FB, AB

