## Supplementary Materials for "Female-dominated disciplines have lower evaluated research quality and funding success rates, for men and women"

Contents:

1. Data Summary table (Table S1)

2. Detailed descriptions of each data set
    - Aotearoa, New Zealand: PBRF research evaluation scores
    - University of Canterbury: bibliometric data
    - Australian Research Council (ARC): funding success rates
    - Canadian Institute of Health Research (CIHR): funding success rates
    - European Institute of Gender Equality (EIGE): funding success rates

3. Electronic data tables details
    - TableE1: Summary statistics of the PBRF data.
    - Table E2: Detailed model output for all PBRF, ARC, CIHR and EIGE data candidate models.
    - Table E3: Detailed model output for the bibliometric models

4. Data files
    - DataARC.xlsx
    - DataCIHR.xlsx
    - DataEIGE.xlsx

### 1. Data summary

| | Scoring research excellence | | Success in research funding | | |
|---|---|---|---|---|---|
| **Country(ies)** | Aotearoa-NZ (PBRF) | | Australia ARC | EU EIGE | |
| **Subnational dataset** | | UC | | | Canada CIHR |
| **Discipline groupings** | 42 (43 in 2018) | 43 | 22 | 8 | 4 |
| **Measures of gender balance** | From national PBRF data (includes all NZ researchers) | | From 2018 survey of Australian Research Population | Gender balance by discipline measured in separate dataset [ref]; or from applicant pool if not available. | From applicant pool |
| **year** | 2006, 2012, 2018 | 2018 (using publications from 2013-18 inclusive) | 2010-2019 (inclusive) | 2019 | Pre-2013 Post-2013 |
| **Other demographic data available?** | Date of birth, Rank (not used) Ethnicity (sometimes, and incomplete) | same | No | No | Original study accounted for researcher age, but unavailable for our study |
| **Does each datapoint represent one individual?** | Yes | Yes | No. represents an applicant; one individual can apply for several grants. | No. Aggregated data reports # applicants, # successes in each country/discipline combination. | No. Aggregated data reports success rates By gender by time point in each discipline? |
| **Sample size** | 2006=4230 2012=6663 2018=7487 | 384 | Over 176,000 applicants aggregated into 440 datapoints (22 disciplines x 10 years x 2 genders) | Over 135,000 applicants aggregated into 333 data points ((8 disciplines x 28 countries x 2 genders) – missing data) | Over 23,000 applicants represented in 16 datapoints (4 disciplines x 2 time points x 2 genders) |
| **Independence** | High within year | same | Some individuals may have applied more than once | Some individuals may have applied more than once | Some individuals may have applied more than once |
| **Quotas/pre-allocation?** | No | No | No pre-allocation of | Unknown | unknown |

| | | | | | |
|---|---|---|---|---|---|
| | | | funds to disciplines before applications scored and ranked. | | |
| **Bibliometric data available?** | No | Per researcher:<br># outputs,<br># weighted outputs (divided by # of co-authors)<br>Mean field weighted citation index<br>Mean SNIP<br>Mean # citations/output | No | No | no |

Table S1: comparative descriptions of each of the 4 datasets

### 2. Detailed data descriptions

**Aotearoa, New Zealand: PBRF research evaluation scores**

New Zealand's Performance Based Research Fund's assessment scores every research-active university academic's research performance from 200-700 points using a personal portfolio. Assessments are holistic and purport to be based more on quality and impact than sheer number of publications (1). A portfolio contains full texts and impact summaries of an academic's nominated four best publications (books, articles, exhibitions, etc.), a list of additional publications and postgraduate supervisions, and evidence of the academic's contribution to the research environment and peer esteem (journal editorial posts, speaking invitations, awards, etc.) during the 6-year assessment period. PBRF takes part-time employment, and special circumstances (illness, heavy administrative load, etc.) into account.

Researchers are assessed by one of fourteen panels consisting of national and international experts grouped by research field; for example, political science and geography disciplines are both grouped under the Social Sciences panel. Panels review their preliminary scores for evidence of patterns of bias before moderating and finalizing the scores.

Assessments give each individual academic three component scores: research outputs, peer esteem, and contribution to the research environment. These are combined in the ratio 70:15:15 to give an overall score from 0 to 700. Scores are then clustered into grades (600-700=A; 400-599=B; 200-399=C; 0-200=R (research inactive)) which are reported back to individuals. Research inactive individuals were not included this study.

| **Time period** | | **2006** | **2007-12** | **2013-18** |
|---|---|---|---|---|
| **Disciplines** | | 42 | 42 | 43 |
| **N** | **Women** | 1708 | 2658 | 3297 |
| | **Men** | 2522 | 4005 | 4181 |
| **Mean Score (std)** | **Women** | 272 (161)$^{***}$ | 368 (139)$^{***}$ | 379 (142)$^{***}$ |
| | **Men** | 351 (163) | 424 (143) | 433 (146) |
| **Mean Age (std)** | **Women** | 44.5 (9.6)$^{*}$ | 47.8 (10.4)$^{***}$ | 47.7 (11.1)$^{***}$ |
| | **Men** | 45.3 (10.1) | 50.0 (10.9) | 49.5 (11.4) |

Table S2: A summary of the three PBRF datasets. Mean (and standard deviation) of score and age at each time point. *** $p < 0.001$, * $0.01 < p < 0.05$, two-sided t-test men/women.

*Discipline groupings:* Individuals self-allocated to one of 42 disciplines covering all fields of research at very high level of detail (see Table E1). At the third assessment in 2018 an additional discipline was added to give a total of 43.

*Gender balance:* As the dataset is the entire population of active researchers the proportion of men in each discipline is taken directly from the data.

*Time:* The dataset covers three time spans (2000-06, 2007-12, 2013-18). We chose to analyse the three time points separately, i.e. treat them as independent datasets. This was due to the relatively long time between assessments and the large number of data points at each time.

*Other available data:* the PBRF dataset also contains detailed data on the research institution of each individual, i.e. one of the eight ANZ universities; researcher date of birth; researcher job title and academic rank, i.e. Professor, Associate Professor etc; researcher ethnicity. In our analysis we accounted for age and institute as continuous and categorical variables respectively. We chose not to include ethnicity as many of the ethnicity groupings were only represented in a small number of disciplines. Job title or academic rank is in itself a measure of research quality albeit a very implicit one with many confounding factors. It has also been shown repeatedly to be subject to many gender biases through promotion and hiring biases. For this reason we did not include it but rather relied on age as a proxy for seniority and experience.

*Sample size:* the final sample sizes for the three sub datasets of PBRF are 2006: 4230 data points; 2012: 6663 data points; 2018: 7487 data points. See Table 1 for the gender breakdown and average research quality of each group.

*Independence:* the three sub-datasets are not strictly independent as many individuals will have been employed at all three assessments. However, at a single assessment all individuals were assessed independently. There were no pre-allocations of scores to different disciplines, i.e. a preponderance of very high scores in one discipline did not require another discipline to be down-graded. Results from the fourteen broad subject area panels were cross-referenced with each other to standardize across disciplines.

*Availability:*

The data used in this study are owned by a third-party organisation (Tertiary Education Commission (TEC), New Zealand). The authors were granted access privileges to the data, under strict nondisclosure agreements, by the TEC for this research project only, under Aotearoa, New Zealand's Official Information Act 1992 which facilitates New Zealanders' access to government records, through a formal information request. All ANZ citizens and residents may make such requests, under the following guidelines: https://www.dia.govt.nz/Official-Information-Act requests. This dataset pertains to thousands of people's employment; hence it is strictly private and highly sensitive. Due to ethical and privacy restrictions, a de-identified data set cannot be made publicly available. Interested researchers are invited to contact the corresponding authors to discuss access to data. Methods Table XX contains summary data by discipline for each time point.

**University of Canterbury (UC) bibliometric data**

We matched the anonymized PBRF data with the UC internal research database using date of birth, gender, ethnicity, discipline, and academic department. Conclusive matches were found for 384 individuals. Researchers with no recorded research outputs were not included. Publications data includes journal publications, books, book chapters and technical reports published during the PBRF assessment period 2012-2018. The publication details of each individual output were combined with information from Scopus about publication outlet, citations (counted in February 2022) and number of co-authors. Where possible, for each individual we calculated the five bibliometrics in Table S7.

*Discipline groupings:* Defined by each individual's 2018 PBRF discipline.

*Gender balance:* The gender balance of the entire ANZ researcher workforce from the full 2018 PBRF dataset.

*Time:* Research score was the 2018 research quality score. Publications were all those recorded between 2013 and 2018, inclusive, the approximate dates of the PBRF assessment.

*Bibliometric data:* the internal database gave a record for every research output by each individual. A research output record included: publication source e.g. journal name, book publisher or report name; Source normalised impact per paper (SNIP – a measure of the journal impact relative to the field); co-author information; citations received by February 2022; field weighted citation impact (FWCI – a measure of the number of citations received with reference to the average number for that field). Research outputs not in journals did not have a SNIP value so were excluded from some calculations. This resulted in five bibliometric variables for each individual. The first two are a measure of the quantity of outputs by a researcher during the assessment period. The final three are more heavily related to the quality of an individual's work.

- $N_{outputs}$ – total number of outputs for that researcher.
- $N_{weighted}$ – total number of outputs weighted by number of co-authors, i.e. a single author publication counts 1, a two author publication counts ½ etc.
- $E(FWCI)$ – mean FWCI across the individual's publications.
- $E(SNIP)$ – mean SNIP across the individual's publications.
- $E(Cites)$ – mean number of citations per publication by that individual.

*Sample size:* 384 individuals were able to be matched with their PBRF data. In total there were 7689 publication records.

*Independence:* Although the dataset did not include the entire research active population of UC researchers there was no bias as the smaller sample was due to matching problems centred around unreliable dates of birth in the two datasets.

*Availability*: Although research publications of staff at the University of Canterbury are publicly available the data linked to the PBRF score of each individual remains confidential as described above.

**Australian Research Council (ARC) funding success data**

Funding success data by gender and field of research for applications to the Australian Research Council are publicly available at https://www.arc.gov.au/gender-outcomes-ncgp-trend-data (Table 4). We used data from 2010 to 2019 on all grant applications by 2-digit Field of Research code as assigned by the applicants. Data included all funding schemes in the National Competitive Grants Programme finalised and announced by the Minister for each year. The number of applicants includes all named researchers on the application, including partner investigators.

*Discipline groupings:* ARC uses 22 Fields of Research that give a relative fine categorization of research discipline. Researchers self-allocated to a discipline grouping.

*Gender balance:* The dataset only contains the gender breakdown of individuals that applied for funding. This is likely to be a biased sample of the researcher population. Instead, we use data from the 2018 Survey of the Australian research population. This gives the number of researchers in each of the 22 disciplines across all Australia universities broken down by gender in 2018.

*Time:* The dataset covers the 10-year period from 2019 to 2019 (inclusive). As the dataset is smaller than the PBRF dataset we choose to treat it as a single dataset with a continuous time variable.

*Other available data:* The 2018 researcher survey also gives a rank breakdown of researchers by gender and discipline. However, as in the case of PBRF we see this variable as being too confounded with grant success to use as an additional predictor variable.

*Sample size:* The data is not at the level of an individual. It is aggregated data giving the number of applicants and the number of successes in each discipline each year. The data is split by gender. In total there are 440 (= 22 disciplines × 10 years × 2 genders) independent data points that include over 176,000 applicants. Note that one individual can apply multiple times in multiple disciplines with varying success, generating multiple 'applicants'.

*Independence:* In the dominant grant categories (Discovery grants) the ARC does not pre-allocate funds to different disciplines during its decision process. Applications are scored and ranked individually to decide funding success.

*Availability:* These are publicly available at https://www.arc.gov.au/gender-outcomes-ncgp-trend-data and https://dataportal.arc.gov.au/era/nationalreport/2018/pages/section1/gender. The subset of the data used in this project are available in Supplementary Material ARCData.xlsx.

**Canadian Institute of Health Research (CIHR) funding success data**

The data are from a previous published analysis by Witteman, Hendricks (8) (Table 2) on the success rates of over 23,000 funding applicants to the CIHR.

*Discipline groupings:* This dataset contains only four disciplines which are all sub-disciplines of medical science (Public Health, Bio-molecular, Clinical and Health Sciences). Researchers self-allocated to a discipline grouping.

*Gender balance:* There was no available data on the gender balance of the researcher population. Instead we used the gender balance of the applicants. As stated previously, this is likely to be biased but unfortunately was the best available proxy.

*Time:* The data is available at two time points: before and after a change in funding processes (i.e. before 2013 there was a single grant category – traditional, this split into foundation and project grants after 2013). The original study found a difference in the success rates of women before and after the change. To account for this we continue to split the data into the two time points and treat this as a categorical variable.

*Other available data:* The original study accounted for researcher age but this aspect of the data is not publicly available. The original study split the post 2013 data into two types of grant application (project and foundation). These were aggregated here as this was not relevant to our study.

*Sample size:* The data is not at the level of the individual. It is aggregated data giving the number of applicants and the number of successes in each discipline at each time point. The data is split by gender. There are 16 (= 4 disciplines × 2 time points × 2 genders) independent data points that include over 23,000 applicants. As with the ARC data one individual can apply multiple times in multiple disciplines with varying success, generating multiple 'applicants'.

*Independence:* No information was available on the independence of the funding allocation. Pre-allocation decisions could have been made to give proportionally higher success rates to some disciplines at a cost to others.

*Availability:* Our study did not use the original dataset described by Witteman (2019). The data used are from analysis of this data and are published in [8] Tables 2 and 3 and summarized in Supplementary Material CIHRData.xlsx.

**European Institute of Gender Equality (EIGE) funding success data**

The data are the funding success rates of Government funded grant applications from the EU and UK from EIGE[28]. Each country is recorded separately.

*Discipline groupings:* The discipline groupings are much coarser than the other datasets. It uses eight disciplines that each cover a broad area e.g. Science, Humanities, Engineering etc. Methods for allocation to particular disciplines are not available.

*Gender balance:* A separate dataset [29] gives the gender breakdown of the same disciplines by country. In a small number of country/discipline combinations there is no population estimate of the gender balance. In these cases the gender balance was estimated using the application gender balance.

*Time:* The dataset is for a single time point 2019.

*Other available data:* There were no other available relevant data.

*Sample size:* The data is not at the level of an individual. It is aggregated data giving the number of applicants and the number of successes in each discipline/country combination. The data is split by gender. In total there are up to 448 (= 8 disciplines × 28 countries × 2 genders) independent data points that include over 135,000 applicants. Note that one individual can apply multiple times in multiple disciplines with varying success, generating multiple 'applicants'. After missing data is removed there are 333 data points. Of the 333 available data points 47 (14%) have no available gender balance and gender balance of the applications is used as a proxy.

*Independence:* No information was available on the independence of the funding allocation. Pre-allocation decisions could have been made to give proportionally higher funds to some disciplines at a cost to others.

*Availability:* These are publicly available at Indicator: Research funding success rate differences (percentage points) between women and men, by field of R&D and in total | Gender Statistics Database | European Institute for Gender Equality (europa.eu) (https://eige.europa.eu/gender-statistics/dgs/indicator/ta_resdig_sctech_funding__sf_fund_appl_ben/datatable) and p_men taken from https://eige.europa.eu/gender-statistics/dgs/indicator/ta_resdig_sctech_rdperes_perf__rd_p_perssci_prop/. Data is collected by EIGE based on the 2021 She Figures publication, based on the women in science (WiS) questionnaire module T1. The subset of the data used in this project are available in Supplementary Material EIGEData.xlsx.

All analysis was done with Matlab (2022b), primarily using the glmfit function.

**Formatted:** Font: Not Bold

### Electronic data tables details

**TableE1: Summary statistics of the PBRF data**

Including the number of women and men, mean age and mean score for each gender and proportion of men in each discipline.

**Table E2: Detailed model output for all PBRF, ARC, CIHR and EIGE data candidate models.**

Each sheet shows the coefficients and p-values for the full range of candidate models trialled for each dataset to predict either Score (PBRF) or Funding success (ARC, CIHR, EIGE). Coefficient names (first column) are given in Wilkinson's notation. Delta AIC is the change in AIC between the best fit model (minimum AIC, Delta AIC = 0) and the candidate model. R-squared is also given for the linear model (PRF only).

Individual country coefficients and p-values for the EIGE data are not shown for clarity.

**Table E3: Detailed model output for the bibliometric models**

Sheet 1: Single bibliometric models $Score{\sim}Age + Gender + \log{(Bib)}$.

The coefficients and p-values for each of the five bibliometrics. Delta AIC is the change in AIC between the best fit model, $N_{outputs}$with minimum AIC and Delta AIC = 0, and the candidate model. R-squared is also given.

Sheet 2: Two bibliometric model $Score{\sim}Age + Gender + \log(N_{outputs}) + \log{(Bib)}$. The coefficients and p-values for each of the other four bibliometrics. Delta AIC is the change in AIC between the $N_{outputs}$ only model (minimum AIC, Delta AIC = 0) and the candidate model. R-squared is also given.

Sheet 3: Coefficients and p-values for $Score{\sim}Age + Gender * p_{men} + \log(N_{outputs})$.

### Data files

**DataARC.xlsx**

Field, Gender, Year, Number of applicants, Number of successes, proportion men (from external data).

**DataCIHR.xlsx**

Field, Gender, Type of grant, Time period (before or after funding change), Number of applicants, number of successes, proportion male (from application numbers).

**DataEIGE.xlsx**

Country, Field, Gender, Number of applicants, Number of successes, proportion male (from applicant data), proportion male from external data.

269
